# miR2Trait: an integrated resource for investigating miRNA-disease associations

**DOI:** 10.1101/2021.11.30.470593

**Authors:** Poornima Babu, Ashok Palaniappan

## Abstract

MicroRNAs are key components of cellular regulatory networks, and breakdown in miRNA function could lead to cascading effects culminating in pathophenotypes. A better understanding of the role of miRNAs in diseases would aid human health. Here, we have devised a method for comprehensively mapping the associations between miRNAs and diseases by merging on a common key between two curated omics databases. The resulting bidirectional resource, miR2Trait, is more detailed than earlier catalogues, uncovers new relationships, and includes analytical utilities to interrogate and extract knowledge from these datasets. The resource could aid in identifying the disease enrichment of a user-given set of miRNAs and analyzing the miRNA profile of a specified diseasome. miR2Trait is available as both a web-server (https://sas.sastra.edu/pymir18) and an open-source command-line interface (https://github.com/miR2Trait) under MIT license for both commercial and non-commercial use. The datasets are available for download at: https://doi.org/10.6084/m9.figshare.8288825.

## INTRODUCTION

MicroRNAs (miRNAs) are key elements of post-transcriptional regulation in the genomic architecture of both prokaryotes and eukaryotes. They are a class of diminutive non-coding RNAs of about 18-25 nucleotides, first observed in *C. elegans* [1]. In miRNA-guided RNA silencing, the miRNA binds to the cognate mRNA and destabilizes it, thereby priming the transcript for degradation. A single miRNA is capable of silencing the expression of several genes by relaxing the specificity of hybridisation. miRNA-based regulation plays crucial roles in health, and miRNA dysregulation is a common mechanism in the aetiology of complex diseases, especially cardio-vascular, auto-immune and neuro-degenerative diseases, and cancers [2-4]. About half of all miRNA genes are present in cancer-related genetic loci [3], and they are known to play key roles in tumorigenesis and cancer progression [5-7].

The naming of human miRNAs in the miRBase registry (https://www.mirbase.org) is related to the miRNA maturation pathway (Figure 1). In brief, miRNA subtyping with numbers (−1, 2, …) indicates identical miRNAs that derive from different genomic regions; miRNA subtyping with letters (a, b, …) indicates nearly identical miRNAs that differ in only one or two positions; and miRNA designations -5p and -3p refer to the source arm of the miRNA duplex from which the mature miRNA is derived. It is the mature miRNA that binds to the mRNA and inhibits protein translation [7].

**Figure 1.**
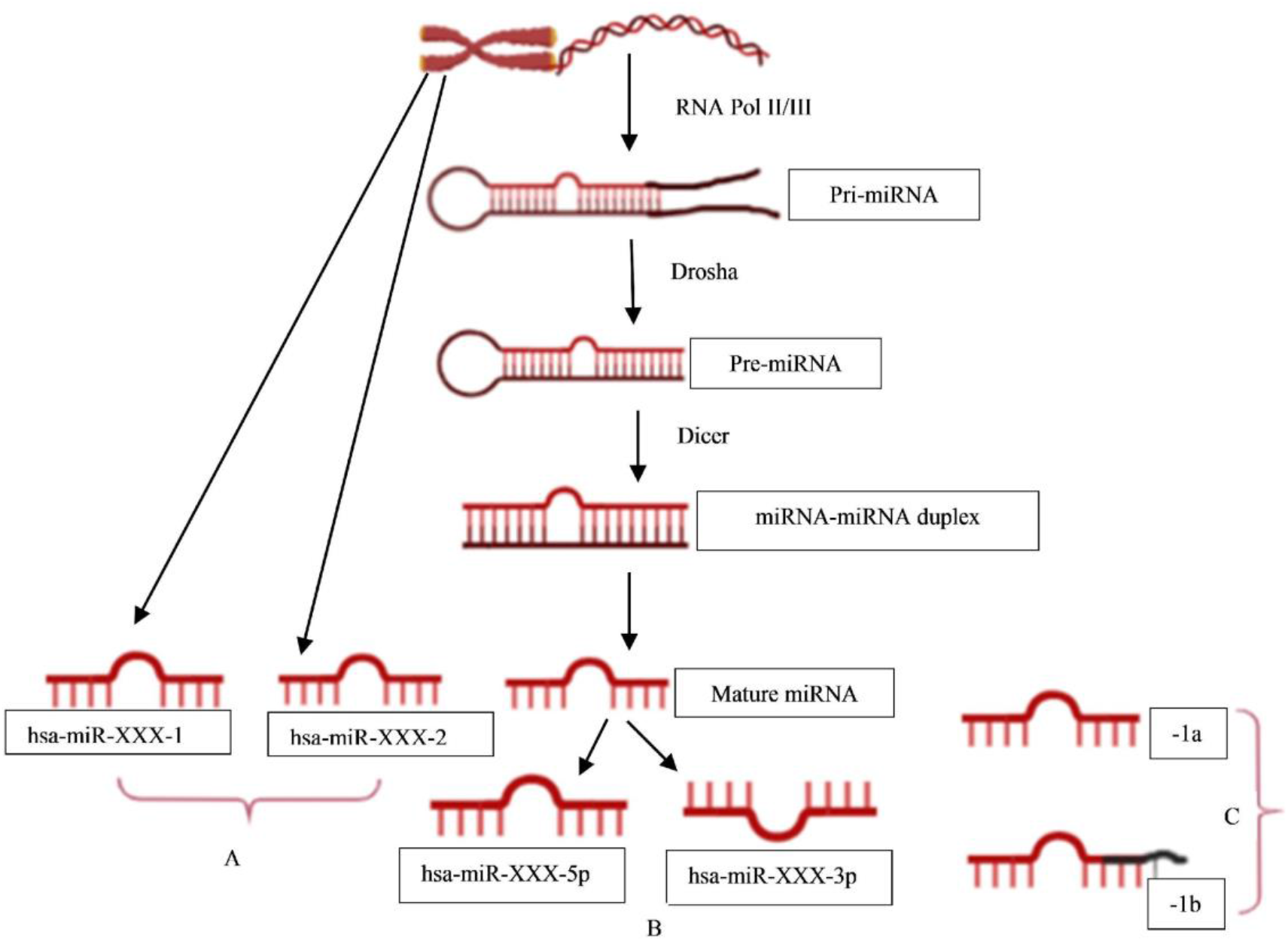
miRNA nomenclature mirrors the pathway of its biogenesis. A: identical miRNAs encoded by different genetic loci (1,2,…); B: source arm of the miRNA duplex (5p or 3p); and C: nearly identical miRNAs that differ in one or two positions (a,b,…).

Existing databases of miRNA–disease associations involve tedious manual curation or are limited in scope and evolution. miR2Disease was one of the earliest catalogues of miRNA-disease associations, providing relationships between 349 miRNAs and 163 diseases [8]. HMDD was developed by Cui et al [9] to provide a more comprehensive database of miRNA-disease associations. It used text mining followed by manual annotation for identifying miRNA-disease connections. The current version of HMDD, version 3.2, contained associations between 1206 miRNAs and 893 diseases, and included miRNA subtype information, but is still missing annotation of the -5p or -3p source arm [10]. Since only one arm of the mature miRNA is predominant *in vivo*, this information could be vital in biological research. In this work, we have addressed the limitations of previous efforts, and designed an algorithm to uncover the spectrum of miRNA–disease associations. Our approach is based on integrating two expert-curated databases on a common key. The resulting resource, miR2Trait, is comprehensive and accurate, uncovers novel relations, and provides assorted tools for investigating miRNA-disease associations. miR2Trait can be queried by both miRNA and disease, and is available as both a web server (https://sas.sastra.edu/pymir18) and command-line resource (https://github.com/miR2Trait/ or http://doi.org/10.5281/zenodo.3342897).

## MATERIALS AND METHODS

### miRNA: disease mapping

Two expert curated databases were used as the source databases in the creation of miR2Trait: (1) miRTarbase, a database of experimentally validated miRNA– target interactions (MTIs), with >13,400 entries [11]; and (2) DisGeNET, a widely used and standardised knowledge management platform with >1,000,000 gene–disease associations [12]. Both databases involved the mapping of genetic information, in one case to miRNAs, and in the other, to diseases. This provided a clue to relating miRNAs and disease via the bridge of genetic information (Figure 2). In-house Python scripts were used to extract the gene:miRNA and the gene:disease mappings from the respective databases as dictionary data structures. The two dictionaries were merged on ‘gene’ to establish a dictionary of miRNA:disease mappings. In a similar vein, an inverse dictionary of disease:miRNA mappings was created.

**Figure 2.**
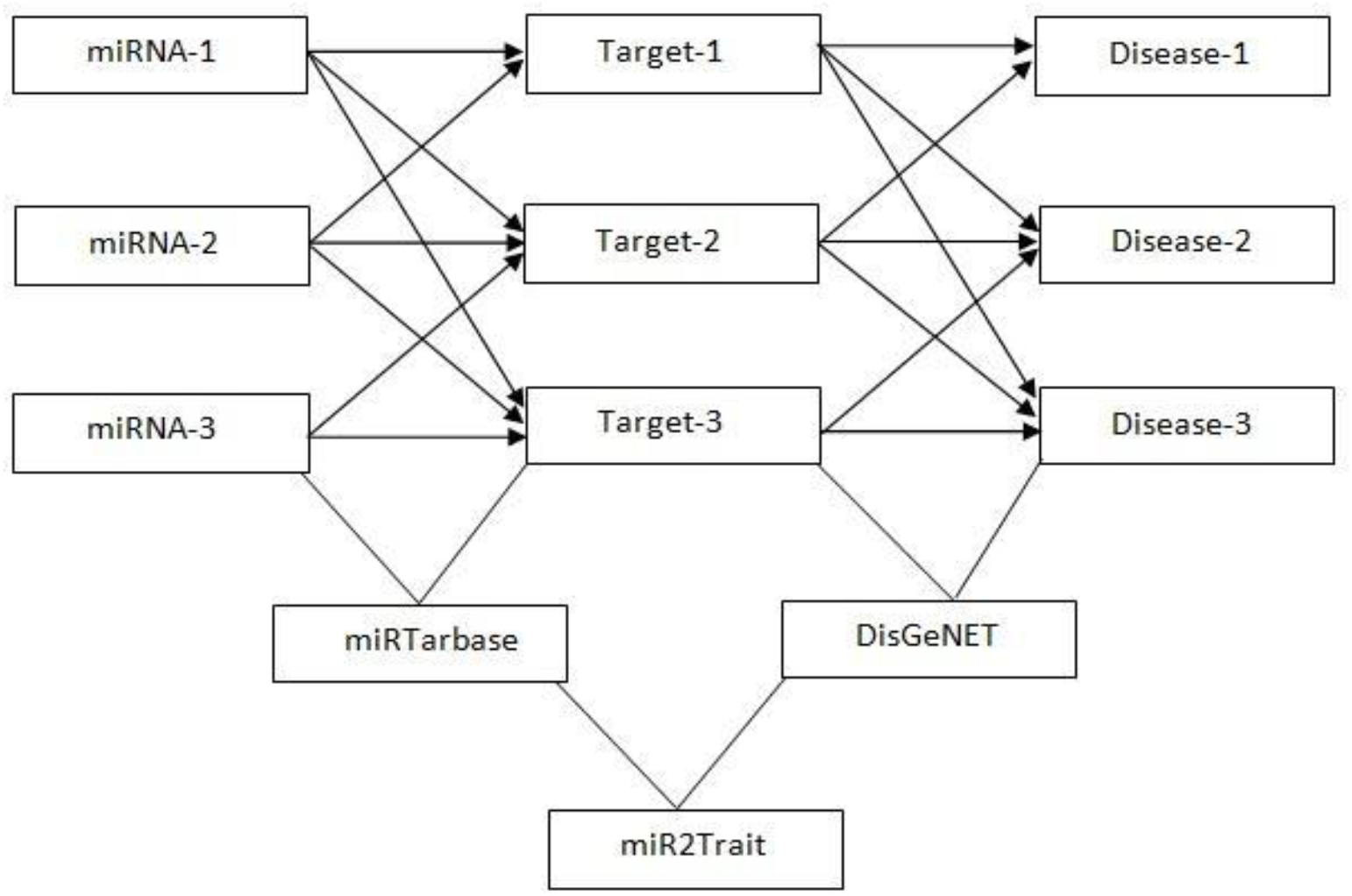
Design of miR2Trait. A back-to-back double bipartite network could be reduced to a single bipartite graph by reductive mapping on the common key. Here, the target nodes of one database serve as the source nodes of the second database. By bridging on these ‘genes’, a mapping between the source nodes of the first database (miRNAs) and the target nodes of the second database (diseases) could be obtained.

### Database creation

The dictionaries obtained above were used to populate relational tables in the construction of two databases: (i) miRNAs and their associated diseases, and (ii) diseases and their associated miRNAs. MySQL was used for encoding the databases.

### Spectrum width calculation

The number of disease associations for a given miRNA is indicative of the breadth of its regulatory impact, which could flag master regulator miRNAs (or regulatory hubs). The disease spectrum width of the i^th^ miRNA is calculated after Qui et al [13] :

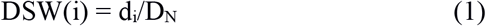

where d_i_ is the number of diseases associated with miRNA i and D_N_ is the total number of diseases in the database. Conversely the number of miRNAs associated with a given disease could be indicative of a complex multifactorial pathology, and the miRNA spectrum width of the jth disease is given by:

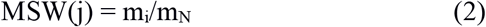

where m_i_ is the number of miRNAs associated with the jth disease and M_N_ is the total number of miRNAs in the database.

### miRNA list enrichment analysis

Given a list of miRNAs that are collectively dys-regulated, it would be of interest to identify which diseases would be enriched (or more likely to occur). Statistical techniques to quantify such enrichment for genes as well as miRNAs exist [14-17]. Here we implement an open-source tool to identify diseases in the miR2Trait database that are enriched for an input set of miRNAs. The method uses a hypergeometric test to quantify the enrichment of each disease in miR2Trait for the given set of miRNAs (in this case identical to the Fisher’s exact test) [18]. The test follows the construction of the Fisher contingency table for each disease, as illustrated in Table 1.

**Table 1.** Fisher’s contingency table for miRNA list enrichment analysis for one disease in the database. Each cell value represents the count for the intersecting row and column.

| miR2Trait | miRNAs associated with disease | miRNAs not associated with disease | (Sum) |
| --- | --- | --- | --- |
| miRNAs in input | a | b | a+b |
| miRNAs not in input | c | d | c+d |
| (Sum) | a+c | b+d | n |

The statistical significance (p-value) of enrichment is computed using Python scipy routines (www.scipy.org), and diseases that pass a user-specified adjusted p-value threshold (with an option to choose among three methods for multiple hypothesis correction) are returned. In addition to significance, both the web server and standalone tool calculate the effect size (log-fold) of each enrichment.

### Diseasome analysis

Given an input set of diseases related in some way (called a diseasome), it would be of interest to find shared dysregulated miRNAs. We approached this problem with an abundance analysis of miRNAs in the diseasome, and identifying miRNAs that passed a user-specified count. Such a set of miRNAs could constitute master regulators acting on common pathways in the specified set of diseases. If we are interested in the miRNA spectra of umbrella disease terms such as ‘carcinoma’, instead of a specific set of diseases, this is also allowed. In this case, diseases containing the general keyword are first identified, and then the miRNA abundance for this diseasome is computed in a manner similar to above.

### miR2Trait web server and standalone tools

A web interface using PHP to connect the user HTML front-end with the MySQL backend was created to offer access to all the services developed, including the two databases and the assorted tools (Fig. 3). MiRNA list enrichment analysis (miLEA) was implemented using the python DBI module, and receives the user’s miRNA list from the HTML form via PHP intermediate. The command-line version of the enrichment analysis provides adjusted p-value calculation using Bonferroni, Holm-Bonferroni or Benjamini-Hochberg (default) correction. All data and code used in the work are available for download, either in the Resources section of the web server or in the github repository.

**Figure 3.**
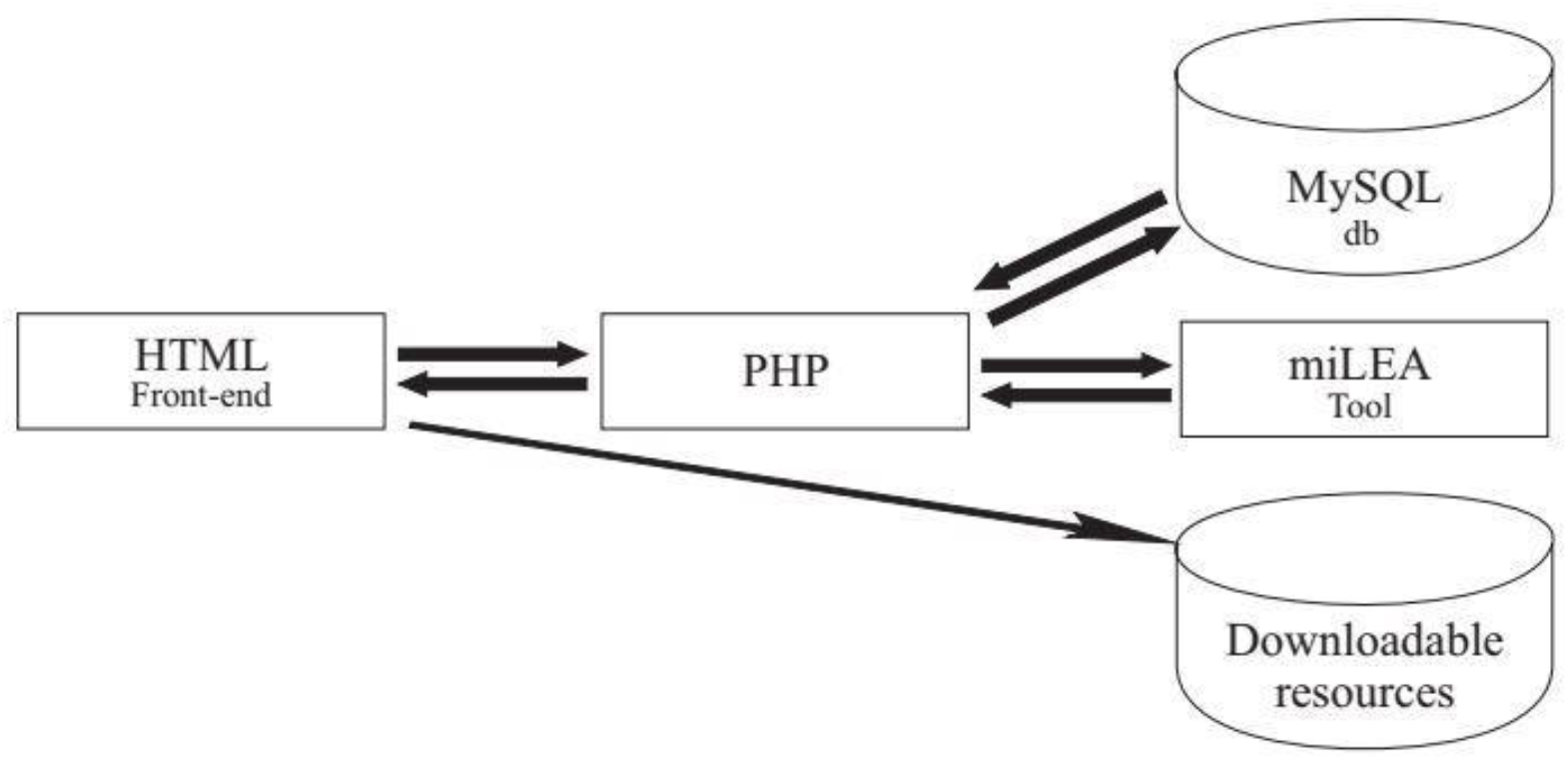
Three layer web-server implementation of miR2Trait accessible at https://sas.sastra.edu/pymir18. The miRNA enrichment analysis backend was implemented in Python.

## RESULTS

The mapping sizes extracted from the respective databases is given in Table 2. In summary, 2599 miRNAs were mapped to 15062 genes using miRTarbase, and 8819 genes were mapped to 13075 diseases using DisGeNET. Merging the two relations yielded a mapping between 2595 miRNAs and 11689 diseases (Table 2). This mapping laid the foundation for the miRNA:disease and disease:miRNA databases, which could be freely downloaded as csv files from aforementioned URLs. The density function of the mappings is shown in Figure 4.

**Table 2.** Establishing the miR2Trait mappings. Only four miRNAs from miRTarBase did not map to any disease in the ultimate analysis.

| Database / resource | Mapping relation | Mapping size |
| --- | --- | --- |
| miRTarBase | miRNAs→Target genes | 2599 → 15062 |
| DisGeNET | Genes → Diseases | 8819 → 13075 |
| miR2Trait | miRNAs→ Diseases | 2595 → 11689 |

**Figure 4.**
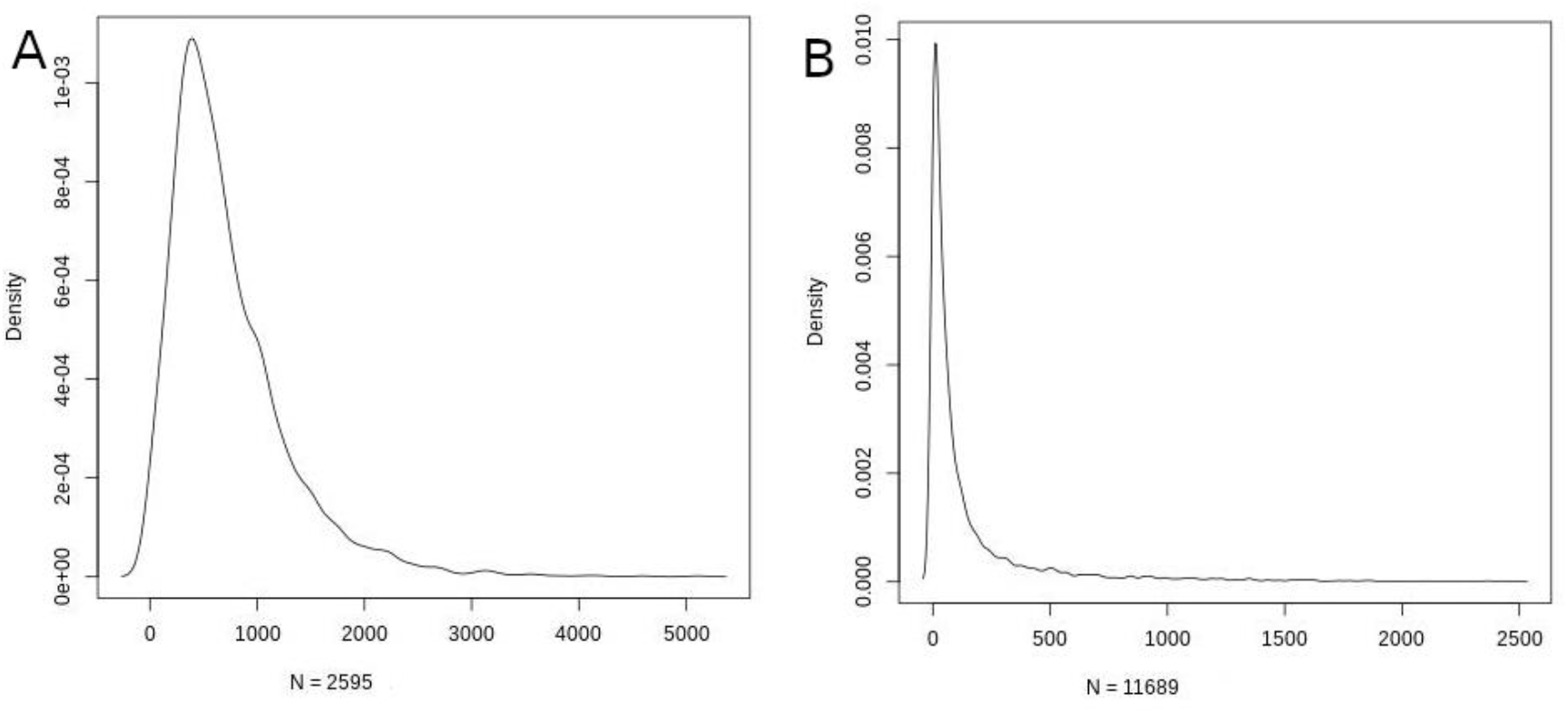
Density function plots of **A)** miRNA-disease; and **B)** disease-miRNA mappings.

Using the miRNA:disease db, miRNAs with the top ten DSWs were identified (Table 3). Such miRNAs are potential master (or hub) regulators at the intersection of multiple pathways, indicatively contributing to one-fifth of all diseases. Using the disease:miRNA db, diseases with the top ten MSWs were identified (Table 4). Complex neurological conditions sweep >90% of all miRNAs, reflecting multifactorial, far from understood aetiologies.

**Table 3.** DSW of miRNAs. miRNA corresponding to the top ten DSWs are shown.

| S.No | miRNA | DSW | No. of associated diseases |
| --- | --- | --- | --- |
| 1 | hsa-miR-335-5p | 0.22 | 2530 |
| 2 | hsa-miR-26b-5p | 0.20 | 2321 |
| 3 | hsa-miR-124-3p | 0.19 | 2175 |
| 4 | hsa-miR-16-5p | 0.18 | 2110 |
| 5 | hsa-miR-92a-3p | 0.17 | 1974 |
| 6 | hsa-miR-1-3p | 0.16 | 1913 |
| 7 | hsa-miR-17-5p | 0.16 | 1912 |
| 8 | hsa-let-7b-5p | 0.16 | 1864 |
| 9 | hsa-miR-155-5p | 0.16 | 1856 |
| 10 | hsa-miR-93-5p | 0.16 | 1766 |

**Table 4.** MSW of diseases. Diseases corresponding to the top ten MSWs are shown.

| S.No | Disease | MSW | No. of associated miRNAs |
| --- | --- | --- | --- |
| 1 | Autosomal recessive predisposition | 0.97 | 2491 |
| 2 | Schizophrenia | 0.96 | 2466 |
| 3 | Intellectual Disability | 0.95 | 2452 |
| 4 | Low intelligence | 0.94 | 2421 |
| 5 | Dull intelligence | 0.94 | 2421 |
| 6 | Poor school performance | 0.94 | 2421 |
| 7 | Mental deficiency | 0.94 | 2421 |
| 8 | Mental Retardation | 0.94 | 2421 |
| 9 | Mental and motor retardation | 0.92 | 2381 |
| 10 | Global developmental delay | 0.92 | 2381 |

The miRNA Enrichment Analyzer computes both the odds-ratio size and significance p-value for a given set of miRNAs. The input could be variable in the number of miRNAs provided. We used “hsa-miR-346 hsa-miR-26a-5p hsa-miR-7-5p hsa-miR-34a-5p” as a query with an adj. p-value (also known as q-value) cutoff of 0.05. This returned a set of 4121 significantly enriched diseases. As another example, we used a set of three random miRNAs: “hsa-miR-2355-3p hsa-miR-214-3p hsa-miR-4801” as a query. This returned 830 significantly enriched diseases (Table 5). Dysregulation of key cancer-associated miRNAs [19] could contribute to the emergence of cancer hallmarks [20] and could be tested likewise.

**Table 5.** miRNA list enrichment analysis. Results for the query: “hsa-miR-2355-3p hsa-miR-214-3p hsa-miR-4801”, and containing the word ‘neoplasm’ are shown, sorted by significance. An odds ratio (effect size) of ‘Inf’ indicates that the entire set of miRNAs is contained in the disease definition.

| Disease | q-value | Odds-ratio |
| --- | --- | --- |
| Colorectal Neoplasms | 0.014 | Inf |
| Neoplasm of the anterior pituitary | 0.020 | 74.50 |
| Stomach Neoplasms | 0.022 | Inf |
| Prostatic Neoplasms | 0.027 | Inf |
| Malignant neoplasm of ovary | 0.034 | 43.32 |
| Biliary Tract Neoplasm | 0.035 | 42.83 |

Analysis of diseasomes for over-represented miRNAs could yield valuable hypotheses for further research, in terms of both pathophysiology and therapeutic options. In this line, we pursued the investigation of diseasomes related to cancer. Since the evidence for miRNA involvement in all stages of cancer including tumorigenesis, progression and metastasis is well-established, we probed miR2Trait for the keyword ‘Neoplasm’ diseasome, and performed an miRNA occurrence analysis. We also performed occurrence analysis for diseasomes identified using the keywords “Carcinoma”, “Sarcoma”, “Lymphoma”, and “Leukaemia”. Table 6 shows the results of these analyses for 25 miRNAs ranked by the ‘Neoplasm’ occurrence analysis. MiRNAs documented in the literature as playing key roles in various cancers appeared in the top miRNAs for each keyword [21-33]. Databases devoted to miRNA associations in cancer have been developed [34-36], and our work here would augment efforts in this direction. Expression of specific microRNAs has been recognized as vital diagnostic / prognostic biomarker and therapeutic target in a variety of cancers [37-45].

**Table 6.**
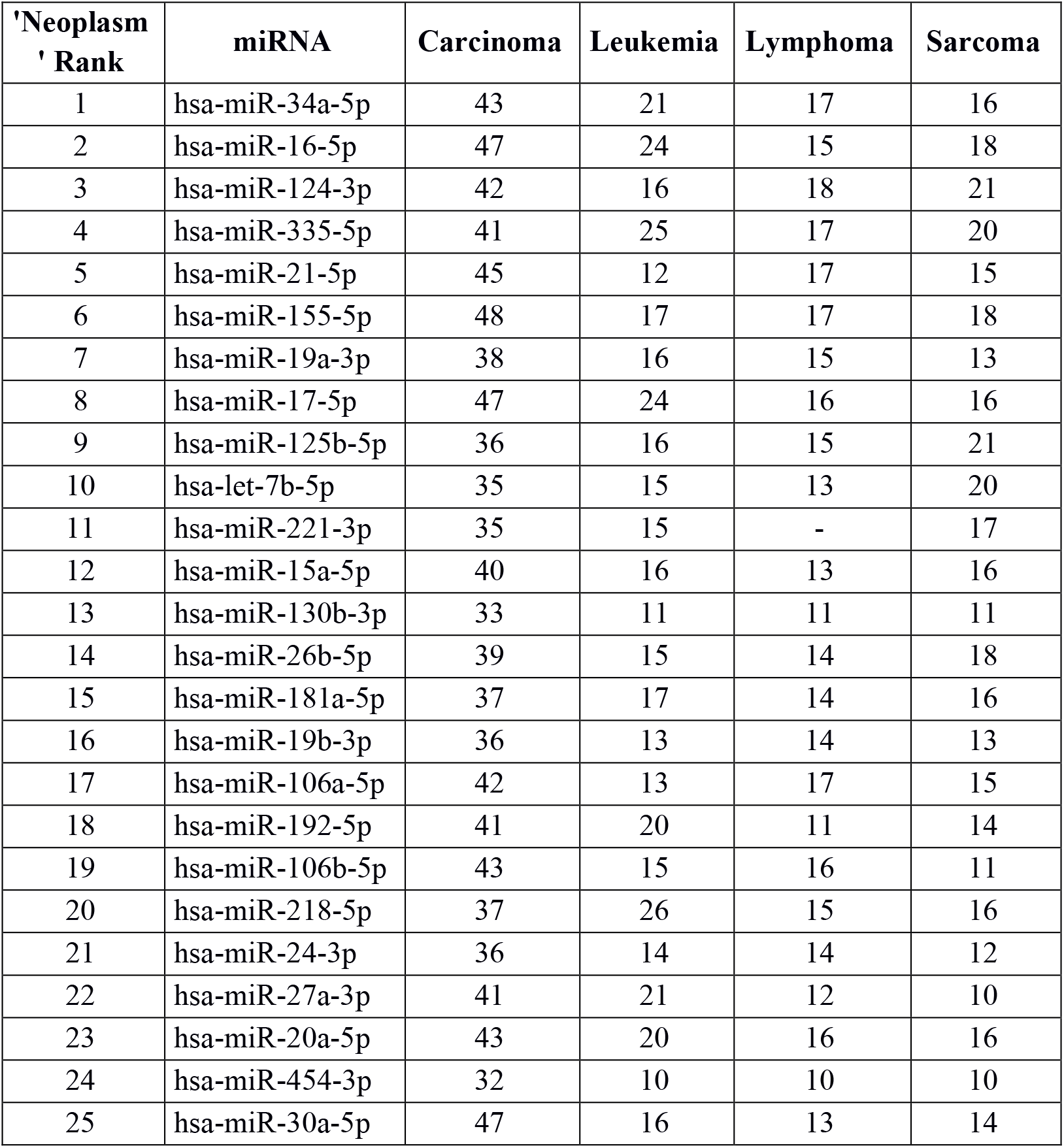
Diseasome analysis of miRNA over-abundance. A summary of cancer-related keyword searches with miR2Trait.

## DISCUSSION

### Benchmarking

Dedicated miRNA omics databases have been developed to aid researchers unravel the role of miRNAs in biological processes (for e.g, see [46, 47]). Computational methods have also been advanced for inferring miRNA-disease connections [48-50]. HMDDv3.2 is a manually curated database of miRNA-disease associations based on text-mining. While miR2Trait contains the full mature miRNA information, HMDD does not provide subtype information such as -5p or -3p; 1a or 1b. The miRNA arm designation (5p or 3p) is a major ingredient of disease aetiology, and key to naming miRNAs (for e.g, the ones with the top DSWs in Table 3). To compare the databases, it is therefore necessary to ‘normalise’ the miRNA nomenclature, by suppressing the subtype information in miR2Trait. This contracted the number of miRNAs in miR2Trait to 1759. Following this, the databases were compared (Figure 5). Even after contraction, miR2Trait appears more comprehensive in the catalog of miRNA-disease associations. MiRNAs unique to miR2Trait showed a 2.5-fold change relative to the miRNAs unique to HMDDv3.2 (884 vs 334). The unique miRNA complements suggest directions for research. There is also a significant overlap between the two databases.

**Figure 5.**
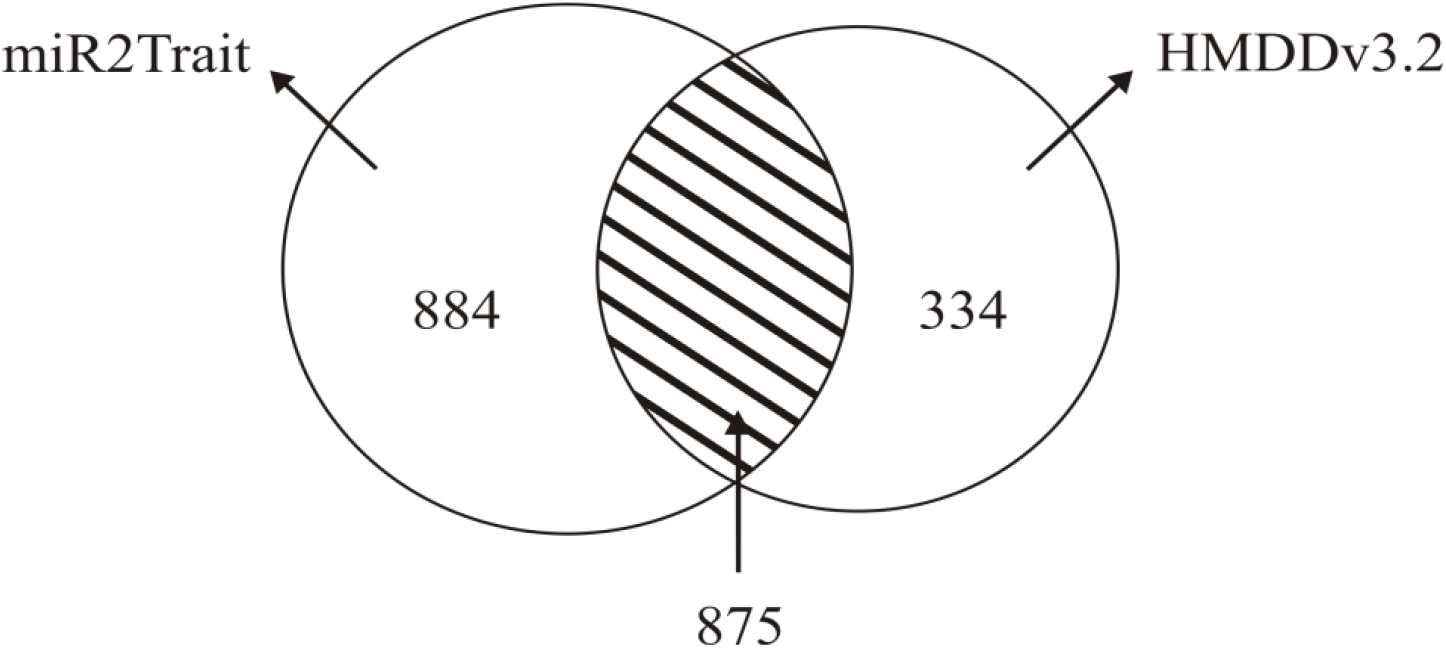
Benchmarking against HMDDv3.2. The analysis revealed a large overlap but also an unexpected asymmetry in the relative sizes of the unique complements. miR2Trait has a clearly larger unique complement.

### Deployment

The miR2Trait web server could be queried by either disease or miRNA. The standard use-case would be to identify miRNAs associated with some disease. For instance, a search by disease for ‘Myocardial infarction’ returned 1445 associated miRNAs. The top hits included hsa-miR-302a-3p, hsa-miR-302b-3p, hsa-miR-302c-3p, hsa-miR-302d-3p, hsa-miR-367-3p and hsa-miR-367-5p, all of which are well-documented in the literature [51]. The top hits for a search by disease of ‘Alzheimer’s disease’ included hsa-miR-9, hsa-miR-29a, hsa-miR34a, hsa-miR-106b, hsa-miR-125b, hsa-miR-146a, and hsa-miR-155, again all of which are well-documented in the literature [52]. As a last example, the top hits for a search by disease of ‘Diabetes mellitus’ included hsa-miR-577, hsa-miR-37a, hsa-miR-375, hsa-miR-181a, hsa-miR-17, and hsa-miR-24, all of which are reported in the literature [53, 54]. Such searches usually also return miRNAs whose links with the query would not be apparent nor readily ascertained from the literature. Search results for such miRNAs lacking documented disease connections might offer hypotheses for future investigations.

The web-server supports advanced searching with regular expressions (https://dev.mysql.com/doc/refman/8.0/en/regexp.html#operator_regexp). The search term is case-insensitive, matches anywhere, and interpreted according to regex grammar. The following use-cases will aid the construction of appropriate queries:

i. To match at the beginning, the ‘^’ symbol may be used. For e.g, ‘^essential.*’ matches diseases beginning with “essential”.
ii. To match at the end, the ‘$’ symbol may be used. For e.g, “.*tension$” matches diseases with “tension” at the end.
iii. To match multiple options, the ‘|’ symbol may be used. For e.g, ‘(cancer)|(neoplasm)’ fetches all ‘cancer’ *and* all ‘neoplasm’ database entries.
iv. To match anywhere, with zero or more occurrences of search term, the ‘*’ wildcard may be used. For e.g., ‘.*tension’ retrieves diseases containing “tension” anywhere in their names; here ‘.’ serves to signify any character. The query ‘hsa-miR-454*,’ when searching by miRNA would match entries that contain “hsa-miR-45” anywhere.
v. The special character ‘+’ mandates the occurrence of the preceding character (at least once). ‘Neo+’ would match diseases containing a prefix word ‘Neo’ (for e.g, any ‘neoplasm’). As another example, when searching by miRNA, ‘hsa-miR-454+’ would match the following: hsa-miR-4540, hsa-miR-454-3p, hsa-miR-454-5p.
vi. A template for constructing queries that retrieve all subtypes of a specific miRNA would be: ‘miR-([0-9][0-9]*)[ab]?-[1-9]-’ where the parenthesis specifies the miRNA of interest. To retrieve only the 5p variants, we could modify the query thus: ‘miR-([0-9][0-9]*)[ab]?-[1-9]-5p’.

## CONCLUSION

MicroRNAs play key roles in the development of disease by virtue of their important regulatory activities. To document such associations, we used a novel method integrating two curated and validated sources of miRNA-gene and gene-disease relations, and created vastly expanded databases of miRNA-disease and disease-miRNA associations. A set of tools to interrogate these databases and uncover novel findings have been developed. Taken together, this work constitutes a resource that would provide evidence for suspected associations, generate new hypotheses for experimental pursuit, and facilitate the investigation of the miRNA basis of disease. A web-server interface to all the functionalities has been developed and is available at https://sas.sastra.edu/pymir18/. The service allows for flexible querying using regular expression-based pattern matching. It includes a tool for miRNA-enrichment analysis that would enable identification of statistically overrepresented diseases in a user-given list of miRNAs. This would help formulate hypotheses of miRNA-mediated common mechanisms underlying multiple disease dysregulation pathologies. A number of examples have been discussed to illustrate the potential uses of the resource. The resource has also been benchmarked with the best alternative miRNA-disease database (HMDDv3.2). Further, the source code for the resource is made available, and includes an additional command-line tool for quantifying miRNA over-abundance in a user-given diseasome. All the data developed in the project are freely available as supplementary information. The resource could potentially contribute to and enhance our understanding of the role of miRNAs in disease.

## ACKNOWLEDGMENTS

We would like to thank the Department of Bioinformatics, School of Chemical & Biotechnology, SASTRA University for infrastructure and computing support. This work was supported in part on DST-SERB grant EMR/2017/000470 to A.P.

## Notes

https://doi.org/10.6084/m9.figshare.8288825

https://github.com/miR2Trait

https://sas.sastra.edu/pymir18

### Competing Interest Statement

The authors have declared no competing interest.

